# A spatial transcriptomic map of subcutaneous adipose tissue aging in male mice

**DOI:** 10.64898/2026.11.24.768963

**Authors:** Hanna Ulrich, Simon Graf, Viktoria Wagner, Jonas Wagner, Nicole Ludwig, Paulina Sprezyna, Thiago Goldbeck-Strieder, Victor G. Puelles, Oliver Hahn, Tony Wyss-Coray, Andreas Keller, Lorenz Adlung, Fabian Kern

**Author notes:** These authors contributed equally (Hanna Ulrich, Simon Graf, Viktoria Wagner). These authors contributed equally.

## Abstract

Aging alters adipose tissue composition and function, but the spatial organization and gene expression dynamics during lifetime remain incompletely characterized. Here we profiled spatial transcriptome-wide changes in the aging male murine inguinal white adipose tissue (iWAT). We describe ten different cell type clusters and three distinct adipocyte states. Further, we detected lipid-associated macrophages (LAMs), the cell type-enriched cluster with the largest number of genes upregulated in old age. LAMs that engage in local crown-like structures (CLS) are associated with pro-inflammatory niche programs at old age while LAMs outside of CLS are enriched in metabolically active transcriptional programmes at young age. LAM spots show reduced predicted lipid oxidation and glycolytic activity with age. Independent validation confirmed functionally reduced glycolytic capacity in aged female and male LAMs. Together, our study provides a deeply sequenced spatiotemporal resource for investigating adipose tissue aging biology, and for prioritizing spatially defined macrophage niches.

## Introduction

Adipose tissue plays an important role in metabolic homeostasis and immune regulation, which is disrupted early during aging^1,2^. Systemic changes in aging converge towards a state of chronic low-grade inflammation, termed inflammaging, also affecting adipose tissue composition and function^3^. White subcutaneous adipose tissue (SAT) has functional implications in energy storage, endocrinology and immune regulation^4^, but undergoes marked age-related changes in morphological and functional properties^2,5^. The advent of technologies, such as single-cell/single-nucleus RNA-sequencing (sc/snRNA-seq), spatial transcriptomics, and 3D tissue engineering, is transforming the field of adipose tissue research^6^. In particular, snRNA-seq protocols have enabled the mapping of white adipocyte heterogeneity in mice and humans^7–9^. Furthermore, integrated spatial transcriptomics data have been used to comprehensively characterize white adipose tissue depots in obese humans^10^. Abdominal subcutaneous adipose tissue from aging humans was profiled by snRNA-seq, and macrophages and mast cells were found to be enriched with age^11^. However, the high-resolution and transcriptome-wide spatial context is missing, as, *e.g*., crown-like structures of macrophages surrounding hypertrophic adipocytes are reduced in adipose tissue after treatment with senolytic drugs in old age^12^. While other organs have been extensively profiled during murine aging^13^, the spatial transcriptomic profiling, in particular the local interplay of parenchymal and immune cells in the adipose tissue during aging remains elusive. We therefore set out to generate a spatial transcriptomics atlas of subcutaneous adipose tissue during aging in 17-, 73-, and 112-week-old male mice, spanning adulthood into advanced age^14^. We reasoned that the subcutaneous depot exhibits high plasticity, thus opening a therapeutic window to maintain or restore physiological properties during aging. Our analyses highlight spatial pattern of adipocyte states and lipid-associated macrophages (LAMs), which we previously characterised as important regulators of metabolic homeostasis in adipose tissue of mice and humans^15^, as populations undergoing age-associated transcriptional changes. We provide evidence that LAMs in aging SAT reduce their transcriptional programmes for glycolysis and oxidative phosphorylation, which we functionally test through Single-Cell Energetic Metabolism by profiling Translation Inhibition (SCENITH)^ref.16^ in an independent cohort of LAMs from SAT of aging mice. As we found the underlying transcriptional changes to be similar in LAMs in the SAT of older human males, our spatial transcriptomics atlas may serve as a resource for the identification of future rational intervention points in adipose tissue aging.

## Results

### Generation of a spatial transcriptomics atlas of murine iWAT aging

Murine inguinal white adipose tissue (iWAT) is a subcutaneous adipose depot. We therefore set out to characterize murine iWAT during aging with 10x Visium spatial transcriptomics profiling (**Fig. 1a**). In total, we analysed iWAT samples from twelve male mice aged 17 (‘young’), 73 (‘middle’) or 112 (‘old’) weeks, also by hematoxylin and eosin (H&E) staining (**Supplementary Fig. 1a**). In the spatial transcriptomics data, we robustly assessed detected features and unique-molecular identifier (UMI) counts for each sample (**Fig. 1b**). Across retained spots, median feature and UMI counts were 9.1-fold and 10.4-fold higher, respectively, than in previously profiled lean murine epididymal adipose tissue^17^, making our dataset a deeply-sequenced and informative resource for the community (**Supplementary Table 1**). Upon strict filtering (*c.f.*, Methods), we ended up with an integrated total of 17,327 quality-control-positive spots (**Fig. 1c**), which we annotated in a semi-supervised manner based on inspection of the H&E stainings and the expression of canonical marker genes (see methods). For instance, we used the macrophage population mMac2, and adipose stem and progenitor cell (ASPC) populations mASPC2/3, which were previously described^7,18^, as a reference for annotation of identified clusters. Furthermore, to visualise LAM-signature enrichment across conditions, we used our published LAM signature^15^ projected on individual spots (**Supplementary Fig. 1b**).

**Figure 1.**
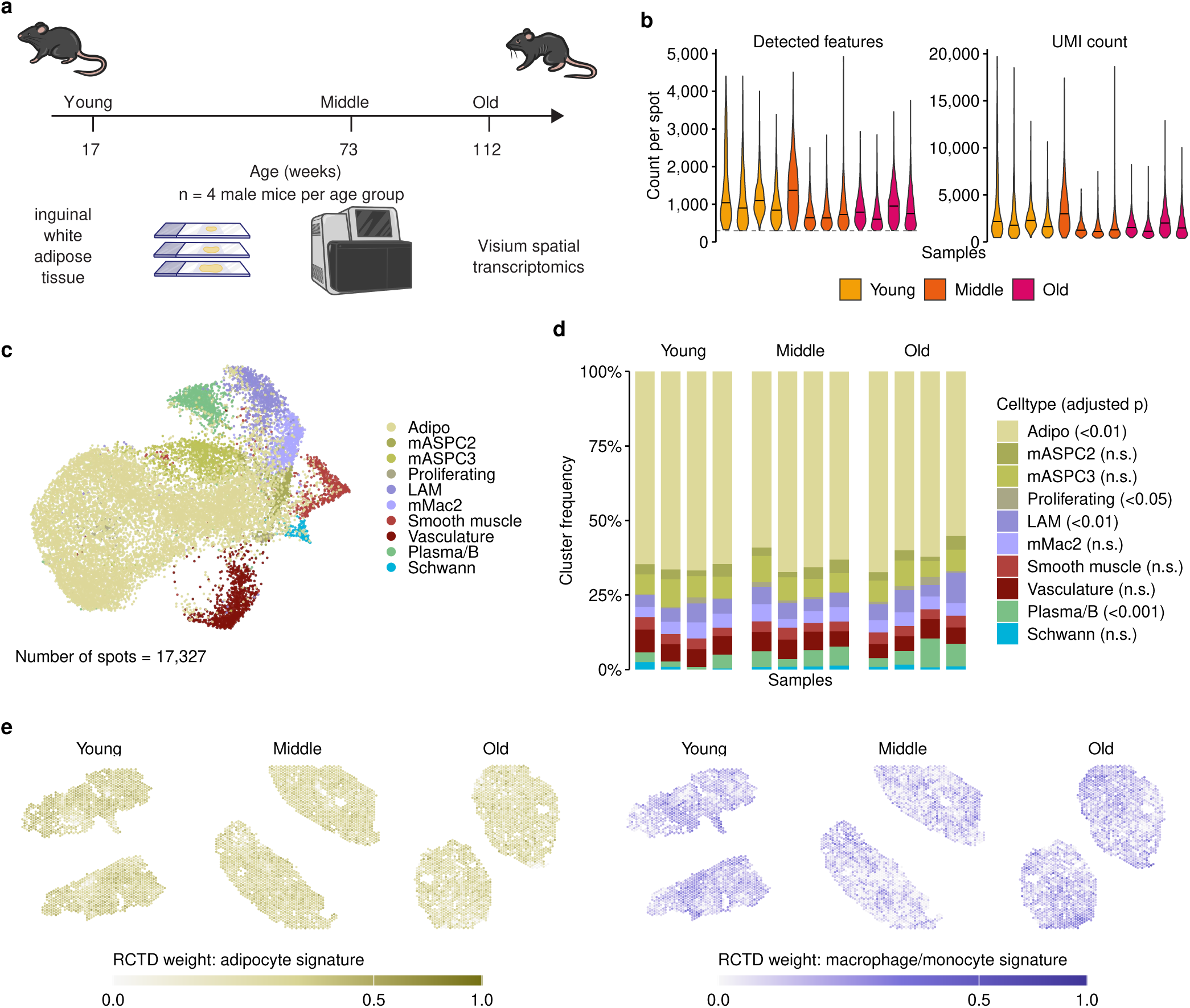
Spatial transcriptomic overview and reference-based compartment mapping of aging inguinal adipose tissue. **a**, Experimental design. Inguinal adipose tissue from mice aged 17, 73 or 112 weeks was profiled by Visium spatial transcriptomics (four biological wildtype samples per age group). **b**, Distributions of detected genes and unique molecular identifier (UMI) counts across spots in each sample; internal lines mark the quartiles, and the dashed line indicates 300 detected genes. **c**, Uniform manifold approximation and projection (UMAP) of 17,327 spots, coloured by annotation. **d**, Annotation frequency in each sample. Values in the legend are Benjamini-Hochberg- adjusted likelihood-ratio p values for the age term in binomial models; n.s., not significant. **e**, Robust cell type decomposition (RCTD) weights for adipocytes and macrophages/monocytes in a representative sample per age group. LAM, lipid-associated macrophage; mASPC, mesenchymal adipose stromal/progenitor cell; mMac2, macrophage population 2.

To explore age-associated differences in observed frequencies of annotated cell-type clusters, we performed spot-level binomial modelling and found adipocyte, LAM, and plasma/B cell annotated spots, beside those with a proliferating signature, to be significantly changing with age (**Fig. 1d**). As spots can contain a mixture of cellular transcriptomes, we sought to corroborate the identity of our annotated spots and performed robust cell type decomposition (RCTD)^19^. This computational method was applied together with a public snRNA-seq dataset (GSE241718) of murine iWAT that contained nine coarse cell clusters^20^. The model-assigned adipocyte weights were distributed broadly across the tissue, whereas macrophage/monocyte weights showed spatially restricted patterns at all three age groups (**Fig. 1e**). For internal quality-control, we compared, within each sample, the median mapped RCTD weight in spots assigned to an annotation with the median in all other spots. Positive median differences across samples supported consistency between the eight mapped annotations and their reference compartments (**Supplementary Fig. 1c**).

These observations underscore that spots with a potential mixture of cellular transcriptomes can be clustered and annotated with dominating cell-types, some of which suggest age-associated changes in frequency in murine male iWAT.

### Resolving adipocyte heterogeneity and cell type-specific age-dependent transcriptomic changes

Then, we set out to characterise our annotated adipocyte spots in more detail. Following sub-clustering of 11,001 spots originally assigned to adipocytes (**Fig. 2a**), we systematically compared the resulting three adipocyte states from our data to the published subsets from Emont *et al*.^7^ Across 1,000 feature-subsampling iterations, adipocyte state 1 mapped most frequently (80.9%) to mAd6 (**Fig. 2b**), which was originally described as an obesity-associated pro-inflammatory, pro-fibrotic adipocyte^7^. Adipocyte state 2 mapped to mAd1 (61.9%) and mAd2 (38.1%), with less frequent mAd1 assignments for state 3 (4.4%) and mAd2 assignments for state 1 (19.1%), collectively suggesting that adipocyte state 2 is a mixture or transition between the other two states identified in our data. Adipocyte state 3 mapped predominantly (95.1% of iterations) to mAd5, which was originally described as a lipogenic adipocyte^7^. To better characterize the functional properties associated with the transcriptomes of our adipocyte states, we performed enrichment analysis of Hallmark pathways among the marker genes of the adipocyte states 1-3 (**Fig. 2c**). We found that adipocyte state 1 was enriched for inflammatory pathways (‘Inflammatory response’, ‘TNF-alpha / NF-kappa-B signaling’), hypoxia and extra-cellular matrix (ECM) remodelling. Adipocyte state 2 only showed a significant enrichment of the MYC target programme, which we consider a rather generic pathway potentially indicative of the transitory nature of this state, or a more immature adipocyte with residual proliferative potential. Adipocyte state 3 was enriched for oxidative phosphorylation, adipogenesis, fatty-acid, xenobiotic and peroxisomal metabolism (**Fig. 2c**). To better understand age-associated changes in adipose composition, we applied our spot-level binomial modelling to the three adipocyte states and found significant changes with a frequency (mean ± standard error of the mean) increase of state 3 (young: 21,71 ± 2,00 %; old: 27,99 ± 1,16 %) and decrease of state 1 (young: 48,85 ± 2,21 %; old: 41,69 ± 2,00 %) at higher age, while state 2 remained unaffected (**Supplementary Fig. 2a**).

**Figure 2.**
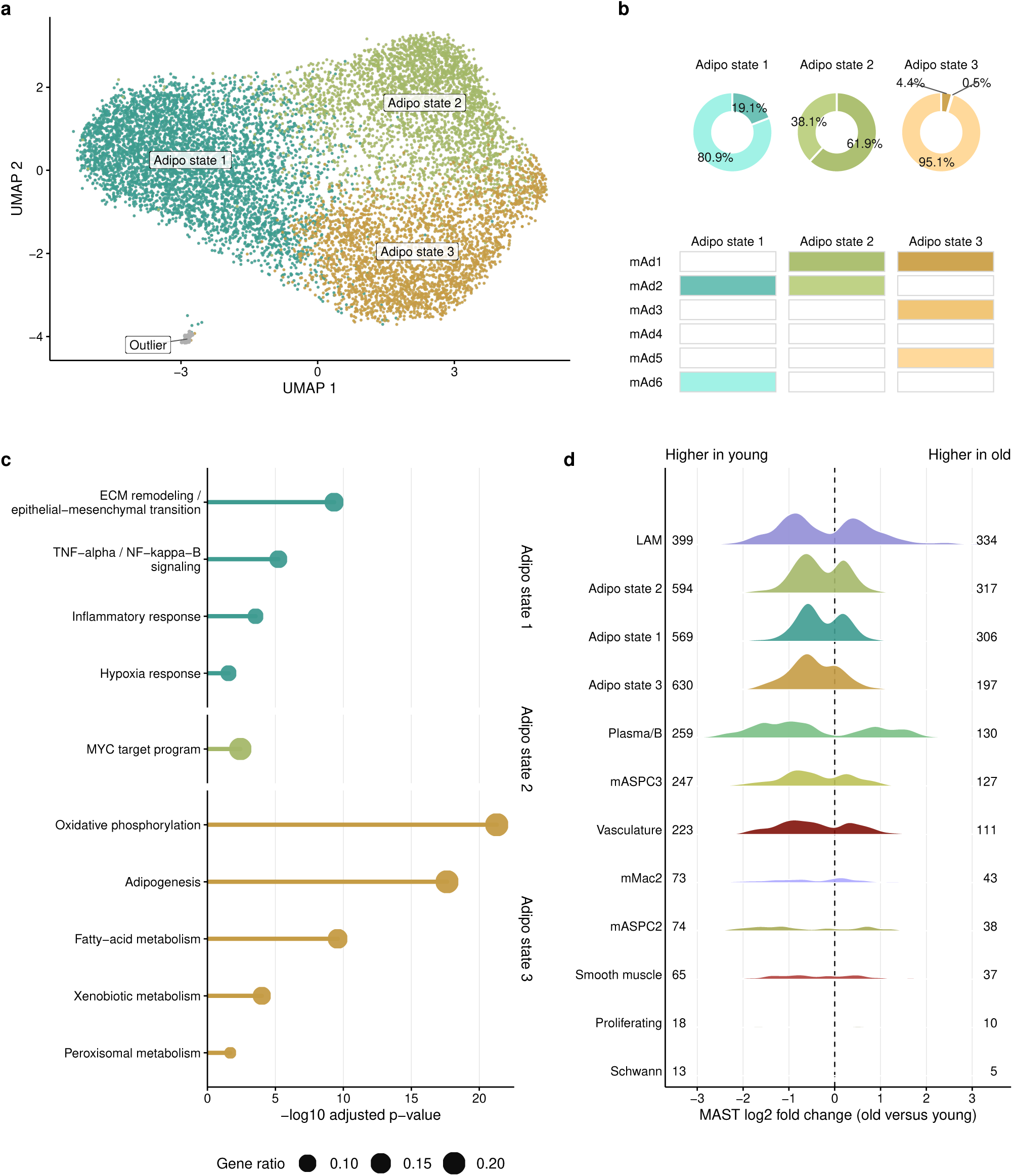
Adipocyte states and age-associated transcriptional changes in inguinal adipose tissue. **a**, Uniform manifold approximation and projection (UMAP) of 11,001 spots assigned to adipocytes, coloured by three robust states; 57 outlier spots are shown in grey. **b**, Mapping stability of each adipocyte state to the mAd1–mAd6 mature-adipocyte populations reported by Emont *et al*.^7^ Donut segments show the fraction of 1,000 feature-subsampling iterations in which each reference population was the top assignment; the matrix shows the observed assignments and their state-matched colour shades. **c**, Selected Hallmark pathways enriched among positive markers of each adipocyte state. Horizontal position denotes the -log10 Benjamini-Hochberg-adjusted p value and circle size denotes the gene ratio. **d**, Distributions of log2 fold changes for genes differentially expressed between 112 and 17 weeks by spot-level model-based analysis of single-cell transcriptomics (MAST). Populations are ordered by the number of upregulated genes and use the population colours from Fig. 1; numbers on the left and right denote downregulated and upregulated genes, respectively. Only genes with Benjamini-Hochberg-adjusted p<0.05 are shown; ridge height is proportional to the total number of displayed genes. ECM, extracellular matrix; LAM, lipid-associated macrophage; mASPC, mesenchymal adipose stromal/progenitor cell; NF-κB, nuclear factor-κB; TNF, tumour necrosis factor. The dataset contains four mice per age group.

Next, we systematically explored the age-associated transcriptional changes within the cell-types of our dataset. To this end, we performed a spot-level model-based analysis of single-cell transcriptomics (MAST)^21^. We reasoned that cell type-enriched genes upregulated with age might be a prime target for anti-(inflamm)aging interventions. When we compared the young and the old mice, we found that LAMs exhibited the highest number (334) of genes significantly upregulated at the old age group (**Fig. 2d**). The three adipocyte states also significantly upregulated many genes in old mice (317 for state 2, 306 for state 1, 197 for state 3), followed by plasma/B cells with 130 and mASPC3s with 127 and vasculature cells with 111 genes. The sets of genes upregulated in old versus young mice overlapped across the three adipocyte states and other cell populations, while some genes showed significant upregulation in only one population (**Supplementary Fig. 2b**). The other two comparisons (middle vs. young and old vs. middle) yielded slightly different distributions (**Supplementary Fig. 2c**), consistent with non-linear age-dependent transcriptional differences and dynamic processes. Complete differential expression results are provided in **Supplementary Table 2**.

### Conserved age-associated transcriptional changes in lipid-associated macrophage tissue niches

Since LAMs exhibited the highest number of genes upregulated in spatially confined tissue niches at old age, we wondered whether there are conserved transcriptional changes of LAMs in aging murine iWAT. To identify concordant transcriptional changes, we performed unsupervised k-means clustering of dynamically changing genes in LAM spots and found five distinct clusters of gene trajectories across the three age groups (**Fig. 3a**). We were again particularly interested in the genes that peak at old age, as those might offer some targets for therapeutic interventions. Cluster 4 contained 4,370 genes that were increasing from 17 to 73 weeks and peaking at 112 weeks of age. When we performed Gene Ontology Molecular-Function (GO-MF) enrichment analysis with the genes from cluster 4, we noted among the top gene sets those associated with ‘GTPase regulator activity’ and ‘nucleoside-triphosphatase regulator activity’ (**Fig. 3b**). To test whether these observed patterns were conserved, we sought to externally validate age-associated LAM transcriptional changes in male murine iWAT. First, we utilized a scRNA-seq dataset of the Tabula Muris Senis^22^. We selected 147 male LAM-signature-enriched macrophages from ten donor IDs (**Supplementary Fig. 3a**) and clustered their gene trajectories across age (**Fig. 3c**). We found cluster 5 with 3,476 genes peaking at old age (96 weeks). When we subjected those genes to GO-MF enrichment analysis, we were able to corroborate ‘GTPase regulator activity’ and ‘nucleoside-triphosphatase regulator activity’ among the top gene sets (**Fig. 3d**). We then repeated our analysis for a published snRNA-seq from the Aging Panorama^23^. We selected 1,176 male LAM nuclei from 16 donor IDs in wild-type iWAT (**Supplementary Fig. 3b**) and clustered their gene trajectories across age (**Fig. 3e**). Cluster 3 contained 3,228 genes peaking at old age (92 weeks). Subsequent GO-MF enrichment analysis with the genes from this cluster yielded again ‘GTPase regulator activity’ and ‘nucleoside-triphosphatase regulator activity’ among the top associated gene sets (**Fig. 3f**). To test to which extent the genes in the clusters peaking at old age were conserved across the three datasets, we inspected the intersection and found 125 genes with a nominal one-sided multi-set intersection p = 1.2e-6 using the 18,176 genes shared by the three datasets as background (**Fig. 3g** and **Supplementary Table 3**).

**Figure 3.**
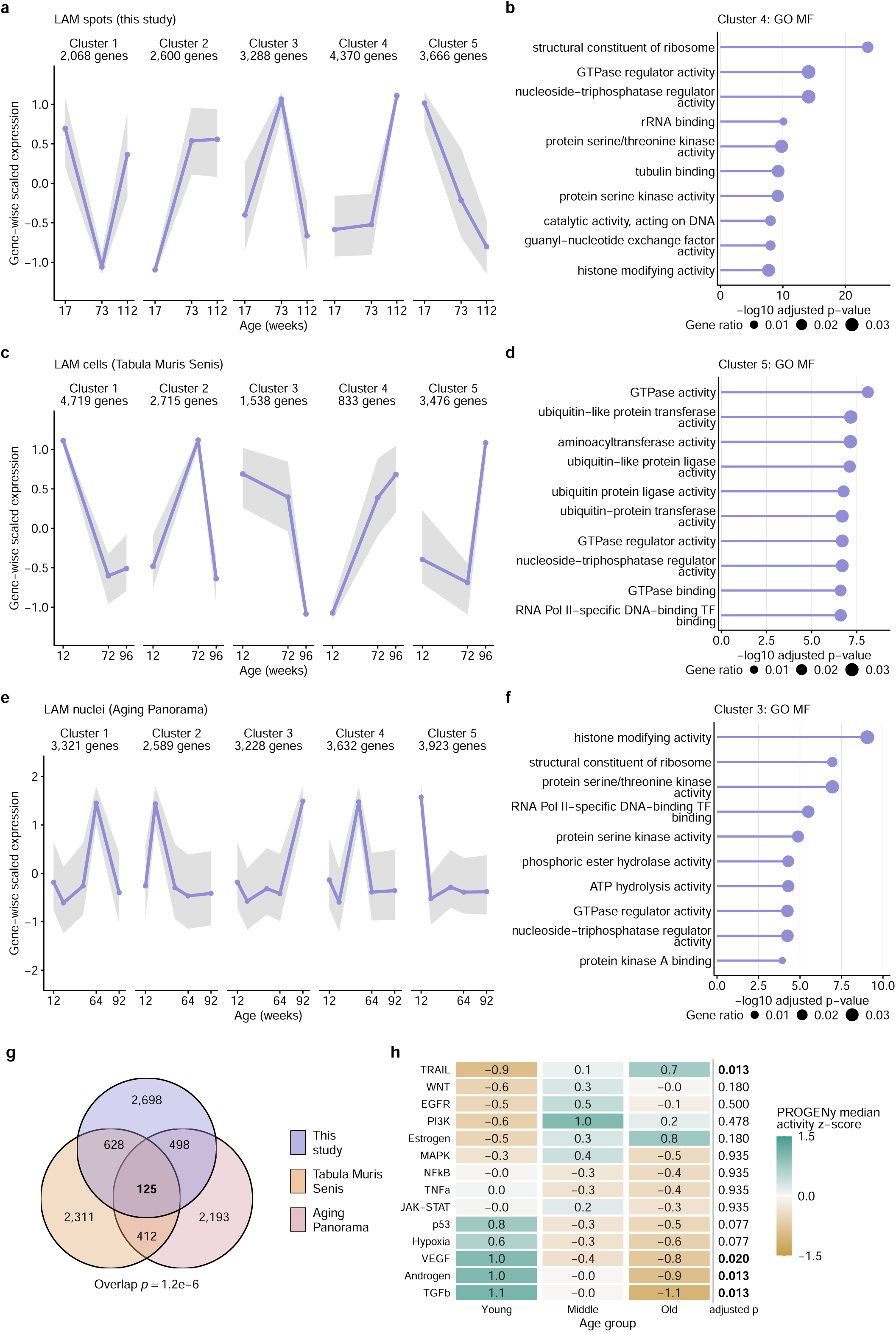
Conserved **age-associated transcriptional programs in lipid-associated macrophages. a**, **c**, **e**, Five k-means clusters of gene-wise scaled mean expression across age in LAM spots from this study (**a**), LAM cells from Tabula Muris Senis^22^ (**c**) and LAM nuclei from Aging Panorama^23^ (**e**). Lines show cluster means and grey bands show the 10th–90th percentile across genes. **b**, **d**, **f**, Gene Ontology molecular-function (GO-MF) enrichment for the late-age-increased cluster in each dataset. **g**, Overlap of the three late-age-increased gene clusters; p denotes the nominal one-sided multi-set intersection test using the intersection of the three input feature catalogues (18,176 genes) as background. **h**, Age-group medians of standardized sample-mean PROGENy^24^ pathway activity in LAM spots at each age, values at right are Benjamini-Hochberg-adjusted p values from two-sided exact age-label permutation tests on sample-mean activities. LAM, lipid-associated macrophage; PROGENy, pathway-responsive gene activity inference.

Since ‘GTPase regulator activity’ suggested altered signal transduction in aging LAMs, we performed PROGENy analysis^24^ and tested sample-mean signalling pathway activities for age-associated changes for all LAM spots in an unbiased, transcriptome-wide manner across the different age groups in our data, displaying age-group medians of standardized sample activities (**Fig. 3h**). TRAIL was the only pathway with a significant positive age trend (adjusted p = 0.013) and is linked to apoptosis induction^25^. Increased inferred TRAIL signalling in aging LAMs raises the possibility of altered apoptotic or efferocytic activity, as TRAIL was shown in other contexts to promote macrophage apoptosis^26^ and enhance macrophage uptake of other apoptotic cells^27^. However, none of the respective gene-expression signatures showed a significant upregulation in aging LAMs (**Supplementary Fig. 3c**). Of note, TGFb signalling was significantly reduced in LAMs with age. Since TGFb signalling in LAMs has been reported to reduce thermogenic functions in adipose tissue^28^, reduction of TGFb signalling-associated transcripts in aging LAMs could be indicative of preserved physiological iWAT functions.

Collectively, our data suggests that aging LAMs in male murine iWAT upregulate transcripts associated with GTPase regulator activity, with concordant enrichments in two published datasets. Since these other data sources do not provide spatial information, we next analysed in detail the LAM neighbourhood and its age-associated changes.

### LAM spatial organisation and metabolic functions with age

Although no significant age trend was detected in LAM phagocytosis-related expression scores (*c.f*., **Supplementary Fig. 3c**), this did not establish unchanged phagocytic capacity. We therefore examined apoptosis-signature scores in non-LAM spots near LAMs in male murine iWAT. We found significantly higher scores in nearby adipocyte-, mASPC3-, vasculature- and plasma/B-annotated spots than in distant spots of the same annotation (**Fig. 4a**). Among the three adipocyte states, LAM-nearby state 1 (pro-inflammatory) and state 2 (transitory) spots showed significantly higher apoptosis-signature scores, whereas state 3 (lipogenic) did not (**Supplementary Fig. 4a**).

**Figure 4.**
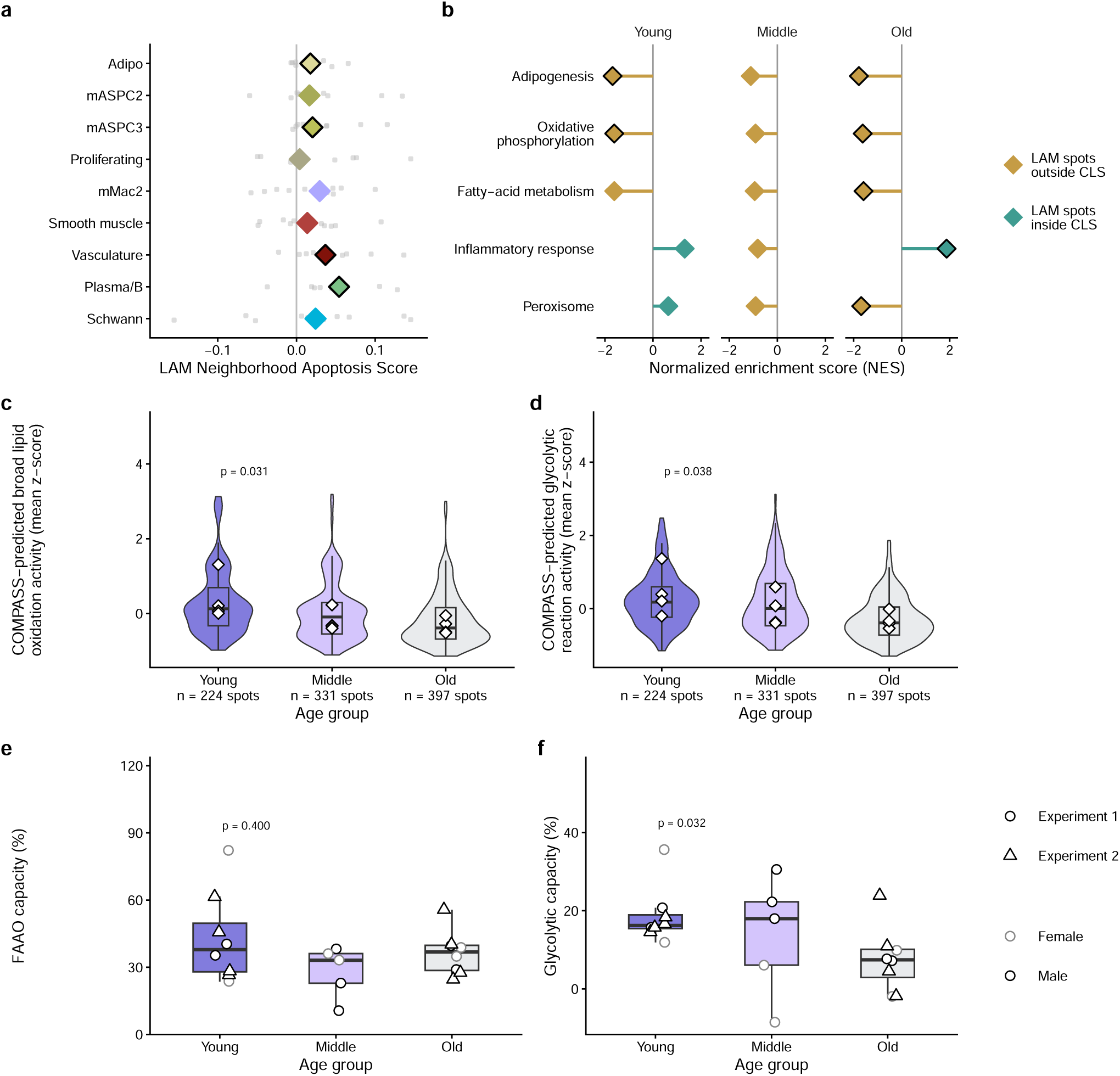
LAM **neighbourhoods and age-associated metabolic remodelling. a**, Difference in apoptosis score between non-LAM spots within 180 µm of a LAM spot and non-neighbouring spots. Grey points show biological samples and diamonds show medians, black outlines denote statistical significance: Benjamini-Hochberg-adjusted p<0.05, exact sign-flip test. **b**, Hallmark gene-set enrichment analysis in paired LAM-spot pseudobulk samples inside versus outside expert-confirmed CLS neighbourhoods. Black outlines indicate Benjamini- Hochberg-adjusted p<0.05. **c**,**d**, Relative COMPASS^31^-predicted broad lipid-oxidation and glycolytic activity, respectively, across all 952 LAM spots. **e**,**f**, SCENITH^16^ combined fatty-acid/amino-acid oxidation (FAAO) capacity and glycolytic capacity, respectively, in LAMs from murine inguinal white adipose tissue. Across two experiment repeats male (young: n=6; middle: n=3; old: n=6) and female cells (young: n=2; middle: n=2; old: n=2) from all age groups were tested. Boxes in **c**–**f** show medians and interquartile ranges; whiskers extend to the most extreme points within 1.5 interquartile ranges. COMPASS violins show spot distributions and white diamonds show mouse medians; SCENITH points represent individual mice. P values calculated from mixed model for age trend with random intercept per mouse and Kenward-Roger degrees of freedom in **c**,**d**, and linear models with sex and experiment id as fixed covariates in **e**,**f**.

Next, we set out to further characterise the spatial organisation of LAMs that surround adipocytes in so-called ‘crown-like structures’ (CLS). Because CLS have been associated with both inflammatory tissue remodelling and macrophage-mediated lipid handling^15,29^, a pathologist manually identified potential candidate CLS in our H&E images. We then performed morphometry of histologically confirmed CLS (**Supplementary Fig. 4b**), which yielded mean diameters of 64.9, 65.2 and 61.8 µm in the young, middle and old age group, respectively (**Supplementary Fig. 4c**). Previous three-dimensional light-sheet microscopy of murine visceral adipose tissue reported mean CLS equivalent spherical diameters of approximately 65 ± 16 µm in lean mice and 74 ± 20 µm in obese mice^30^.

To test whether there are gene expression changes within the local niches of CLS, we performed a gene-set enrichment analysis (GSEA) for Hallmark pathways of the transcriptomes of LAMs inside or outside CLS, respectively (**Fig. 4b**). We found that pathways associated with adipogenesis, oxidative phosphorylation, fatty-acid metabolism and the peroxisome were significantly enriched in LAM spot transcriptomes outside of CLS at the young or the old age group. On the other hand, the inflammatory response was significantly enriched in LAM spot transcriptomes inside our expert-annotated CLS in old mice (**Fig. 4b**). This argues for distinct transcriptional profiles of LAMs inside and outside of CLS. These results suggest spatially distinct metabolic and inflammatory expression programmes to play a role in aging iWAT.

To better understand, based on their spot transcriptomes, how metabolic functions of LAMs change with age, we utilized COMPASS^31^. This framework integrates gene expression with a genome-scale metabolic network to infer relative metabolic reaction activities. Since we found oxidative phosphorylation to be enriched in LAM spots outside CLS (**Fig. 4b**), we assessed age-associated changes in broad lipid oxidation across all 952 LAM spots from all twelve mice. We found that the COMPASS-predicted broad lipid oxidation activity significantly decreased in LAM spots with age (**Fig. 4c**). To assess whether another central metabolic programme showed a similar age association, we also calculated the COMPASS-predicted glycolytic reaction activity and found it to be significantly decreased in LAM spots with age (**Fig. 4d**).

### Functional and human validation of LAM transcriptional changes with age

Next, we sought to functionally validate those transcriptome-based predictions with the SCENITH technology^16^. To this end, we applied the flow cytometry-based approach to analyse male and female LAMs from murine iWAT within the three different age groups (young: 13-18 weeks (n=8); middle: 67-77 weeks (n=5); old: 95-119 weeks (n=8)), allowing us to profile their combined fatty acid and amino acid oxidation (FAAO) capacity and glycolytic capacity (**Supplementary Fig. 4d**). Mean FAAO capacity decreased from 43% to 28% from the young to the middle age group and slightly recovered to 36% in old mice (**Fig. 4e**). On the other hand, mean glycolytic capacity significantly decreased from 19% to 14% to 8% between the young, middle and old age group (**Fig. 4f**). These findings support the transcriptomic trends that aging LAMs in murine iWAT lose to varying extent their capacity of oxidative phosphorylation and glycolysis.

To test the translational relevance of our findings on aging LAMs, we re-analysed two independent publicly available datasets of human aging subcutaneous adipose tissue. We re-analysed GSE175495^ref.3^^2^, a bulk RNA-sequencing dataset of a total of eleven younger and eleven older male donors drawn from a study cohort with reported mean ages of 27 and 66 years, respectively. We performed GSEA of Hallmark gene sets associated with oxidative phosphorylation and glycolysis, and we found a significant enrichment of oxidative phosphorylation in younger samples, whereas glycolysis showed a nonsignificant trend in the same direction (**Fig. 5a**). As glycolysis was not significantly enriched in this dataset, we assessed whether the bulk transcriptomes contained an age-associated LAM-signature enrichment. Our LAM signature^15^ was significantly enriched in the older samples similarly to the Hallmark gene set associated with an inflammatory response (**Fig. 5b**).

**Figure 5.**
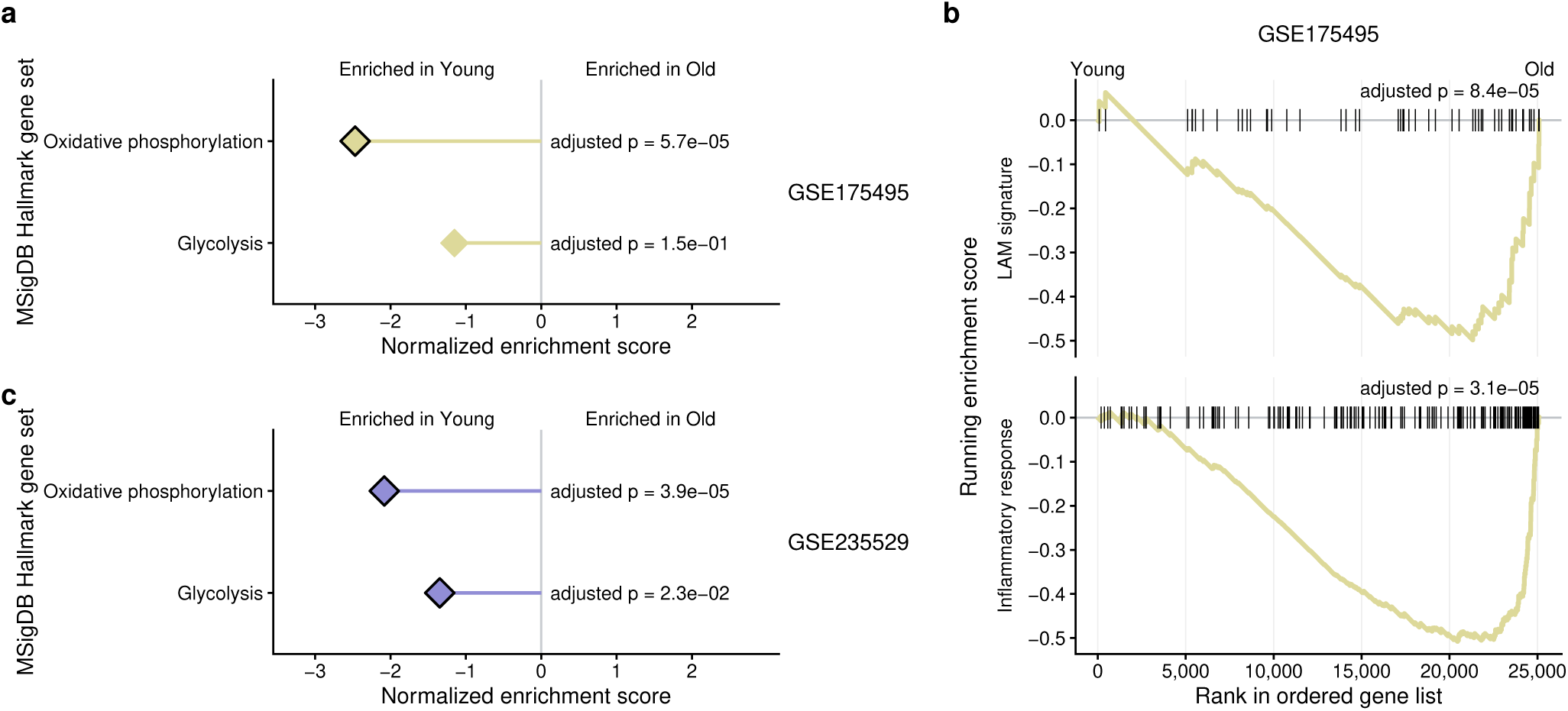
Human **transcriptional support for age-associated adipose metabolic remodelling. a**, Gene-set enrichment analysis of Hallmark oxidative phosphorylation and glycolysis in GSE175495 subcutaneous adipose bulk RNA-sequencing from eleven younger and eleven older men. **c**, Corresponding enrichment analysis in GSE235529 LAM-nucleus donor pseudobulks from five younger and five older men, with BMI adjustment. Black diamond outlines mark Benjamini-Hochberg-adjusted p<0.05; exact adjusted p-values are given. **b**, Running enrichment curves for the LAM signature^15^ and the MSigDB Hallmark inflammatory-response gene set in GSE175495. Black ticks mark contributing genes. Benjamini-Hochberg-adjusted p-values are given.

To identify LAM-specific gene expression changes in human aging subcutaneous adipose tissue, we utilized GSE235529^ref.11^, a snRNA-seq dataset with a total of five younger and five older male donors from a cohort with reported mean ages of 26 and 74 years, respectively. We first identified a cluster of LAM nuclei (**Supplementary Fig. 5a**). Of note, there was no significant difference in the fraction of recovered LAM nuclei between the age groups (**Supplementary Fig. 5b**). We then performed GSEA of MsigDB^33^ Hallmark gene sets associated with oxidative phosphorylation or glycolysis. Both pathways were significantly enriched in LAM nuclear transcriptomes from younger male human subcutaneous adipose tissue (**Fig. 5c**).

Together, these results support that LAMs in aging subcutaneous adipose tissue of humans and mice adapt their transcriptome towards less glycolytic and respiratory pathways potentially influencing parenchymal cells in their surrounding spatial tissue niches.

## Discussion

Adipose tissue is increasingly recognised as immune-modulating depot in aging research but remained incompletely characterized in the spatiotemporal domain. In a single coherent cohort of wildtype male mice, we characterize heterogeneity in observed adipocyte states and LAMs in the context of iWAT aging. Limitations of the first generation Visium technology prevent an annotation at the single-cell level, as a single spot potentially involves a mini-bulk ensemble of different cells. This calls for future validation through subcellularly-resolved spatial transcriptomics technology like Visium HD. There have been attempts to integrate single-cell/nucleus and spatial transcriptomics information on adipose tissue in mice and humans comparing adipose tissue physiology^34^, aiming towards a consensus annotation^35^. We contribute to resolving existing uncertainty by providing coherent high-quality tissues in a sample size superseding those of previous reports^17^ combined with manual expert H&E stain screenings.

Our results point at reduced lipid oxidation and glycolytic capacity in aged LAMs that correlate with a spatially confined imprint into the surrounding neighbour cells, especially in native crown-like tissue structures, or niches. The ‘metabolic exhaustion’ of aged LAMs is consistent with previous findings that aging compromises macrophage bioenergetics through impaired de novo NAD⁺ synthesis and dysregulated glucose utilization^36,37^. Mechanistically, reduced Ca^2+^ uptake into mitochondria was linked to metabolic dysfunction and inflammaging in peripheral macrophages^38^. Aging adipose tissue macrophages were reported to suppress adipocyte lipolysis through NLRP3-dependent catecholamine degradation^39^, pointing to a loss of tissue-remodelling LAM functions and concomitant acquisition of pro-inflammatory properties in aging.

The somewhat weaker effects observed in our SCENITH validation *ex vivo* may partly reflect metabolic adaptation to the assay conditions^16^. When glucose becomes limited, macrophages were shown to increase the contribution of glutamine to the TCA cycle^40^. However, in human aging, macrophages have been reported to concomitantly lose utilization capacity for alternative substrates, including glutamine^37^. Compensatory amino acid metabolism therefore remains an untested possibility in aged LAMs isolated from murine iWAT, particularly because standard SCENITH does not distinguish fatty acid from amino acid oxidation capacity^16^.

Even though our spatial transcriptomics cohort consisted of male mice only, our validation studies on the SCENITH experiment and independent experiments from the literature suggest that metabolic impairment may also occur in female macrophages. Nevertheless, sex can modify the adipose tissue response to aging, as aged obese female mice exhibit a lower accumulation of pro-inflammatory CD11c⁺ adipose tissue macrophages than age-matched males with similar body weight^41^. Thus, sex-specific differences in LAM aging warrant direct investigation.

In the future we aim to broaden the spatial scope to other molecule types, including proteins, metabolites and bioactive molecules such as cytokines, to obtain a more comprehensive picture on the LAM cell state and function within aged versus young native tissue. This will help us to narrow down potentially localized cell-to-cell communication regimes that actively shape the development of the crown-like structures, prompting also their role in aging. To this end, we provide our entire results as interactive and free-to-use online resource to the scientific community under: <u>adlunglab.com/GeroLAM</u>.

## Methods

### Mice

Mice bred in the UKE animal facility were of C57/BL6J wild-type background. Animals were kept in accordance with the institutional review board ‘Behörde für Soziales, Familie, Gesundheit und Verbraucherschutz’ (Hamburg, Germany), and under specific-pathogen-free conditions at ambient temperature. The spatial transcriptomics cohort comprised male mice aged 17, 73 and 112 weeks (four mice per age group) as previously described^13^.

### Tissue processing and spatial transcriptomics

Procedures on tissue collection, Visium chemistry, sectioning, permeabilization, library preparation and sequencing were performed as previously described^13^. The optimal permeabilization time for adipose tissue was optimized and determined to be 15 minutes at a slice thickness of 16 μm.

### Bioinformatics preprocessing and sample integration

Imaging and sequencing data preprocessing as well as sample integration, clustering, filtering, quality control and annotation was performed as previously described^13^.

### Spatial annotation and adipocyte states

Reference-based deconvolution used RCTD^19^ (spacexr, full mode) and an inguinal adipose single-nucleus reference from GSE241718^ref.20^, retaining nuclei with at least 200 genes and 500 UMIs and at most 20% mitochondrial counts. Reference clusters were annotated by canonical markers. Within each mouse, median RCTD weights in each mapped annotation were compared with those in other spots. Spatial LAM-signature enrichment was scored using AUCell^42^. Adipo spots were reclustered in Seurat^43^ (v5.4.0) using 30 principal components and Leiden resolution 0.3, yielding three states and a 57-spot outlier group excluded from state- level comparisons. State markers were tested using sample-blocked edgeR^44^ pseudobulks. Mapping to published mAd1–mAd6 populations used pseudobulk expression similarity and 1,000 iterations sampling 80% of shared genes; percentages represent mapping stability. Hallmark enrichment used positive state markers (Benjamini Hochberg (BH)-adjusted p < 0.05; log2 fold-change ≥ 0.25), with tested genes as background. Age-variable genes were ranked by variance of age-group mean.

### Age-associated differential expression

Pairwise age contrasts within spatial populations used MAST^45^ with log-transformed detected-feature counts as a covariate. Genes expressed in at least 10% of spots in either comparison group were tested; P values were BH-adjusted within population and comparison. Significant genes were displayed without a fold-change cutoff. Sample-level edgeR pseudobulks provided a sensitivity analysis. To visualize overlapping gene sets that were significantly upregulated in old versus young mice per annotated cell-type, we utilized an UpSet plot^46^.

### Reference LAM selection and cross-dataset comparisons

Myeloid populations in Tabula Muris Senis^22^ and Aging Panorama^23^ were ranked by median expression of the published 59-gene LAM signature^15^ relative to expression-matched control genes. Clustering and scoring used both sexes; temporal analyses used males only. The displayed populations were Tabula Muris cluster 0 (Louvain resolution 0.4; 147 cells from 10 donor IDs) and published Aging Panorama cluster 10 (1,176 nuclei from 16 donor IDs). Age-group mean expression was standardized per gene and partitioned into five k-means clusters. Genes from clusters increased at old age were compared by gene-ontology (GO)^47,48^ Molecular Function (MF) over-representation analysis with clusterProfiler^49^. The full org.Mm.eg.db gene universe with the GO-MF-annotated genes served as effective background. BH adjustment was performed within each enrichment run. The three-way gene overlap was tested by an exact one-sided multi-set intersection test, with all sets restricted to the 18,176 Ensembl^50^ genes, which were shared among the three gene lists, as background.

### Pathway scores and spatial neighbourhoods

PROGENy^24^ activities were inferred with each 500 footprint genes per pathway. Scores for Hallmark apoptosis, phagocytosis/engulfment and apoptotic-cell-clearance were calculated from normalized expression. Age trends were tested by exact two-sided age-label permutations of mean scores at mouse level. P values were BH-adjusted across all 14 PROGENy pathways; or jointly Holm-adjusted across the three LAM process scores. For neighbourhood analysis, non-LAM spots within 180 µm of a LAM spot were compared per mouse with same-annotation spots that were more distant. Paired differences in mean apoptosis score were tested by exact two-sided sign-flip tests. P values were BH adjusted either across all nine annotations, or separately across the three adipocyte states. Age dependence of paired differences was tested by an exact age-label permutation test using an omnibus ANOVA F statistic. P values were BH adjusted across all nine annotations.

### Crown-like structures and metabolic predictions

A human pathologist reviewed all 227 H&E candidate structures, classifying 44 positively as CLS, 110 as negative and 73 as indeterminate. For CLS morphometry, we reviewed outer contours of immediately surrounding cellular crowns that enclosed the central adipocyte area. Circle-equivalent diameters were calculated as means of mouse-specific medians of 2√(contour area/π) from the H&E images. CLS neighbourhoods comprised spots within 150 µm of confirmed CLS, but excluded indeterminate or overlapping regions. LAM counts were analysed as paired inside/outside pseudobulks through sample-blocked edgeR quasi- likelihood models with age-specific region contrasts. Results were ranked by signed square-root F statistics for fgsea Hallmark enrichment. P values were BH adjusted. COMPASS analysis^31^ (Mouse1 model) was applied to all 952 LAM spots. Reaction scores of broad lipid-oxidation and glycolytic activities were transformed and normalised. Age trends were calculated using restricted-maximum-likelihood mixed models with a mouse random intercept and Kenward-Roger degrees of freedom.

### SCENITH

Mice were anaesthetized with an 80:20 CO₂/O₂ mixture and sacrificed by cervical dislocation. iWAT from both sides was collected and pooled from two mice when the total tissue weight was below 0.2 g. Tissue was transferred into phenol red-free DMEM (Thermo Fisher Scientific) and kept cool. After mincing, tissue was digested in 5 mL collagenase II solution (Sigma-Aldrich) at 37°C for 15 min under constant agitation. The resulting cell suspension was sequentially passed through a metal strainer and a 250-µm filter, followed by centrifugation at 350 × g for 5 min. The adipocyte fraction was removed, and the stromal vascular fraction (SVF) was treated with 1x ACK red cell lysis buffer (Thermo Fisher Scientific) for 5 min. The SVF was washed, resuspended in PBS and seeded into a 96-well plate.

SCENITH was performed according to previously published protocols^16,51^, with minor modifications. After equilibration for 30 min at 37°C and 5% CO₂, cells were treated with 2-deoxy-D-glucose (2-DG, 100 mM, Sigma-Aldrich), oligomycin (0.1 µM, Cayman Chemical), or both inhibitors (2-DGO). For combined treatment, 2-DG was added first, followed by oligomycin 5 min later. After 15 min of inhibitor treatment, puromycin (20 µg/mL, Sigma-Aldrich) was added for 30 min. The plate was then placed on ice, and cells were washed with cold FACS buffer (PBS supplemented with 2% FCS). Surface staining was performed in 50 µL FACS buffer using anti-CD11b FITC (clone M1/70, BioLegend, 101205), anti-CD45 BUV395 (clone 30-F11, BD Biosciences, 564279), anti-CD63 PE-Cy7 (clone NVG-2 BioLegend, 143909), anti-CD9 PE (clone MZ3, BioLegend, 124805) and anti-F4/80 BV421 (clone BM8, BioLegend, 123137) antibodies, together with a Live/Dead stain (Zombie Aqua™ Fixable Viability Kit, BioLegend) for 20 min at 4°C. Cells were washed in FACS buffer and fixed and permeabilised using the True-Nuclear™ Transcription Factor Buffer Set (BioLegend) according to the manufacturer’s instructions. For intracellular staining, cells were incubated in 25 µL permeabilisation buffer supplemented with 20% FBS for 10 min at room temperature, followed by overnight incubation with anti-puromycin AF647 (clone 2A4, BioLegend, 381508) at 4°C. Finally, cells were washed, resuspended in FACS buffer and analyzed by flow cytometry (BD LSR II, BD Biosciences). Combined fatty acid and amino acid oxidation capacity and glycolytic capacity were calculated as described elsewhere^16^.

### Human transcriptomic analyses

GSE175495^ref.32^ bulk adipose RNA-seq counts were filtered with filterByExpr, TMM-normalized and analysed using limma-voom^52^. For GSE235529^ref.11^, pseudobulks of LAM nuclei were calculated per donor. Models included age group and, for GSE235529, body mass index (BMI). Moderated t statistics were used to rank genes for fgsea enrichment with 100,000 permutations. LAM nuclear fractions in GSE235529^ref.11^ were compared using a two-sided exact permutation test at the donor level.

## Statistical analysis

Statistical tests and multiple-testing correction were conducted in R (v4.4.0) and specified for each analysis. Biological replication refers to mice or human donors. Exact tests, pseudo-bulk models and mixed models were used as described in the analysis-specific methods and figure captions.

Age-dependent spot frequencies were analysed using binomial generalized linear models with categorical age. Likelihood-ratio tests compared models with and without the age term; P values were Benjamini-Hochberg-adjusted across ten main annotations or three adipocyte states, respectively. Computational workflows, parameters and software versions are provided in the accompanying code on <u>github.com/AdlungLab/GeroLAM</u>.

## Data availability

The sequencing datasets and images generated throughout this study are available in the NCBI Sequence Read Archive (SRA) via accession-ID PRJNA1561417 as well as Gene Expression Omnibus (GEO) via accession­ID GSE354520. Summary statistics for individual analyses are provided in the supplementary information files. Spatial transcriptomic data generated in this study will be made publicly available upon publication through the Universität Hamburg Research Data Repository at doi.org/10.25592/uhhfdm.22091. Previously published datasets analysed in this study include the mouse adipose tissue snRNA-seq reference dataset GSE241718; the mouse adipocyte reference from Emont *et al*. (GSE176171); the Tabula Muris Senis scRNA-seq data (GSE149590); the inguinal white adipose tissue subset of the Aging Panorama snRNA-seq atlas (GSE247719); and the human subcutaneous adipose tissue bulk and snRNA-seq datasets GSE175495 and GSE235529, respectively.

## Code availability

The code and required data sources to reproduce all main findings of this work will be made freely available upon peer-review of this manuscript under: github.com/AdlungLab/GeroLAM.

## Statement on the use of generative artificial intelligence tools

We follow a ‘human-first & human-last’ strategy when it comes to the use of generative artificial intelligence (AI) tools and large language models (LLMs). This means we will never prompt tasks that we could not do ourselves, and we critically review all output generated by those tools so that we take full responsibility of all claims made. For coding and writing assistance, we used GPT-6 Astra.

## Acknowledgements

This study was funded by the M.J. Fox Foundation (MJFF-021418), the Phil and Penny Knight Initiative for Brain Resilience, the Schaller-Nikolich Foundation, the Hans-und-Ruth-Giessen Foundation, Saarland University and University of Hamburg. Computational resources used within this study were financed through the Deutsche Forschungsgemeinschaft (DFG) (project no. 469073465). JW received support from the DFG (project no. 493624519). L.A. acknowledges support from the DFG (project no. 528292361) and from the Daimler and Benz Foundation in cooperation with the Reinhard Frank Foundation (project no. 32-01/24).

## Author contributions

H.U. performed spatial transcriptomic and flow cytometric experiments, analysed the data and revised the manuscript. S.G. implemented and performed computational workflows.

V.W. performed spatial transcriptomic experiments and supported data analysis. J.W. identified crown-like structures. N.L. supported sample preparation, spatial transcriptomic experiments and sequencing. P.S. assisted sample preparation. T.G-S. and V.G.P. assisted quality control of the spatial transcriptomic samples. O.H. contributed to study design, mouse cohort generation and sample extraction. T.W-C. and A.K. supported study design, analysis and interpretation. F.K. and L.A. conceived the study, analysed the data, drafted the manuscript, and revised the manuscript. All authors approved the manuscript.

