## Supplemental Data 1 for "A spatial transcriptomic map of subcutaneous adipose tissue aging in male mice"

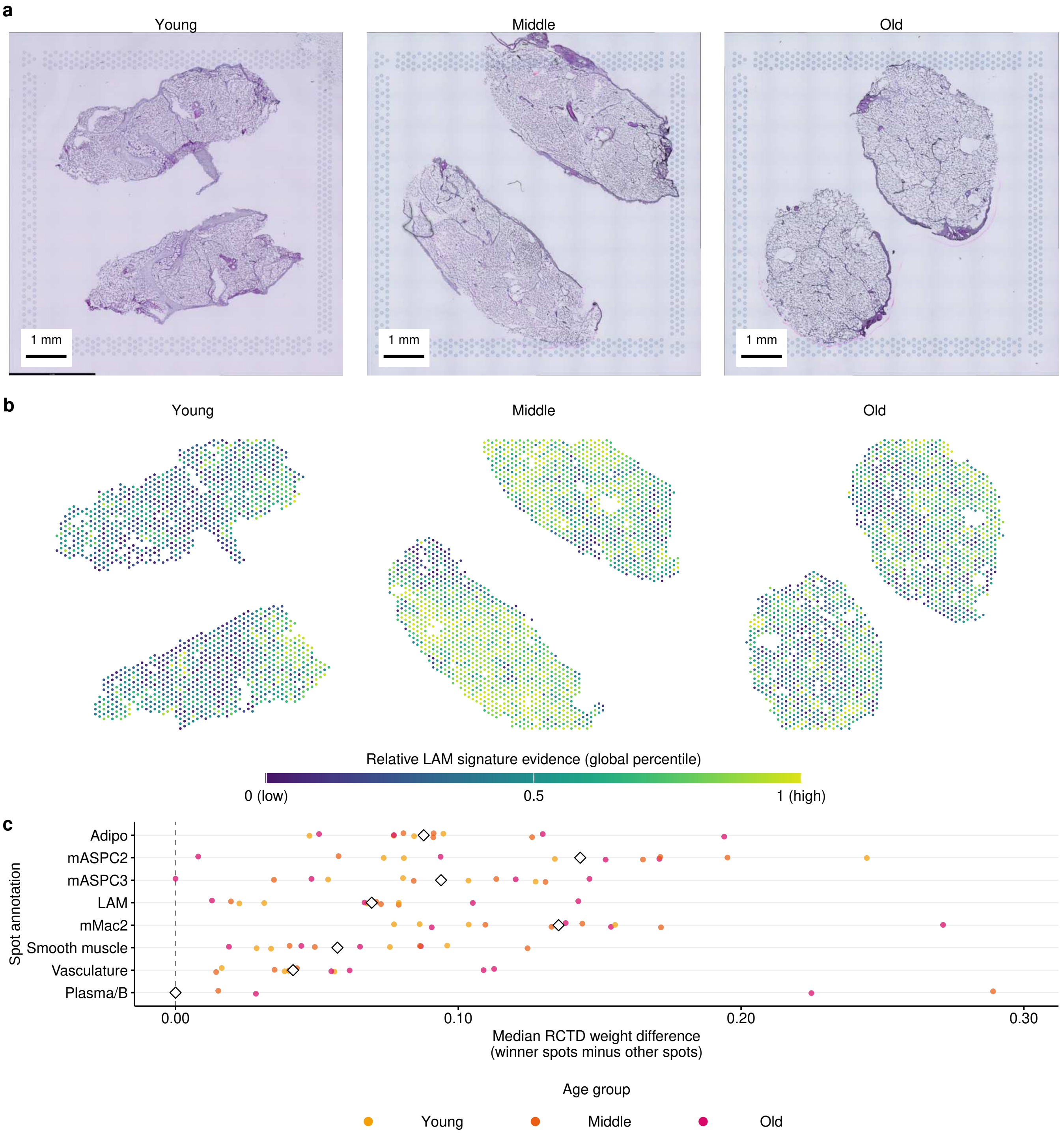


**Supplementary Figure 1**

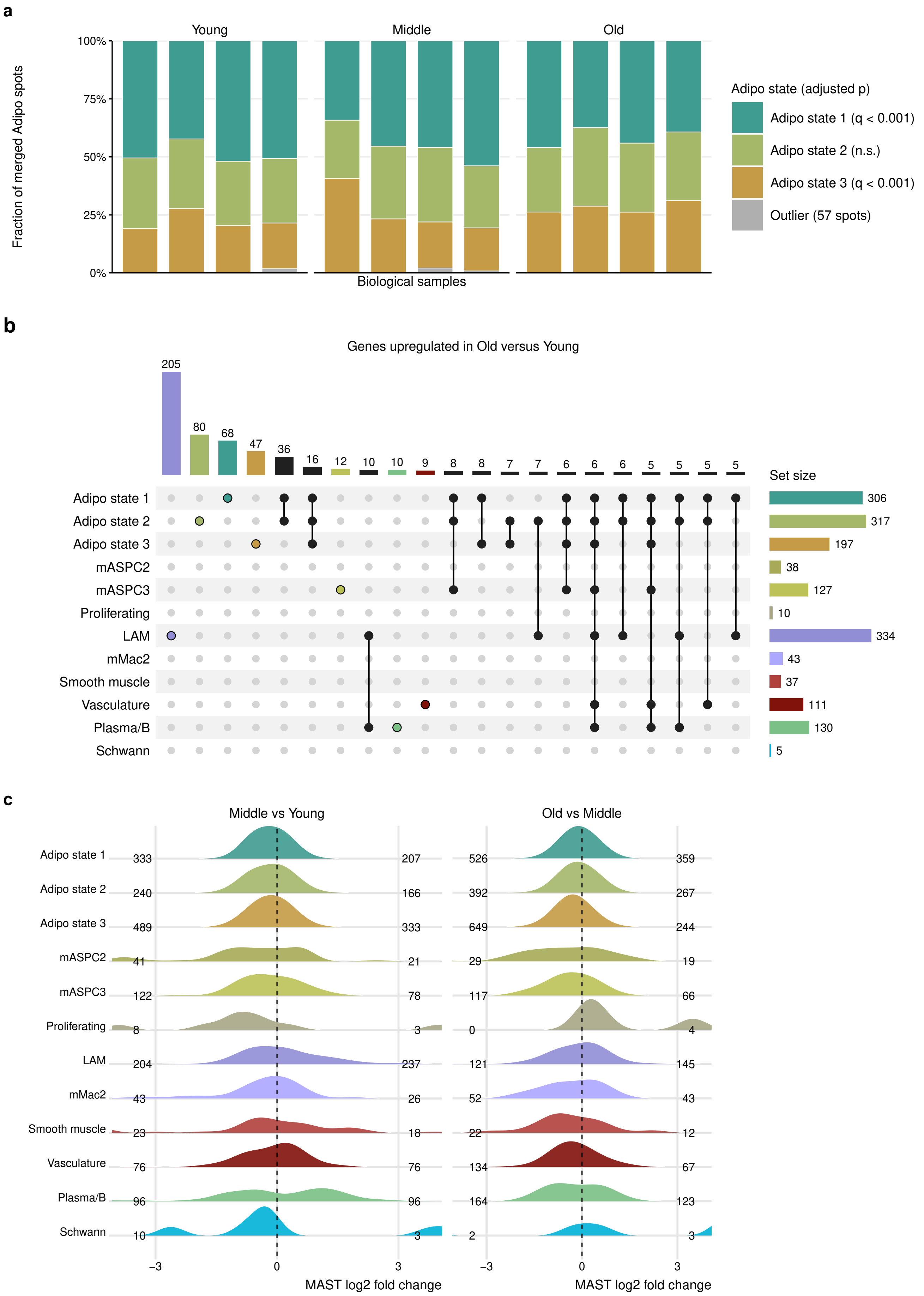


**Supplementary Figure 2**

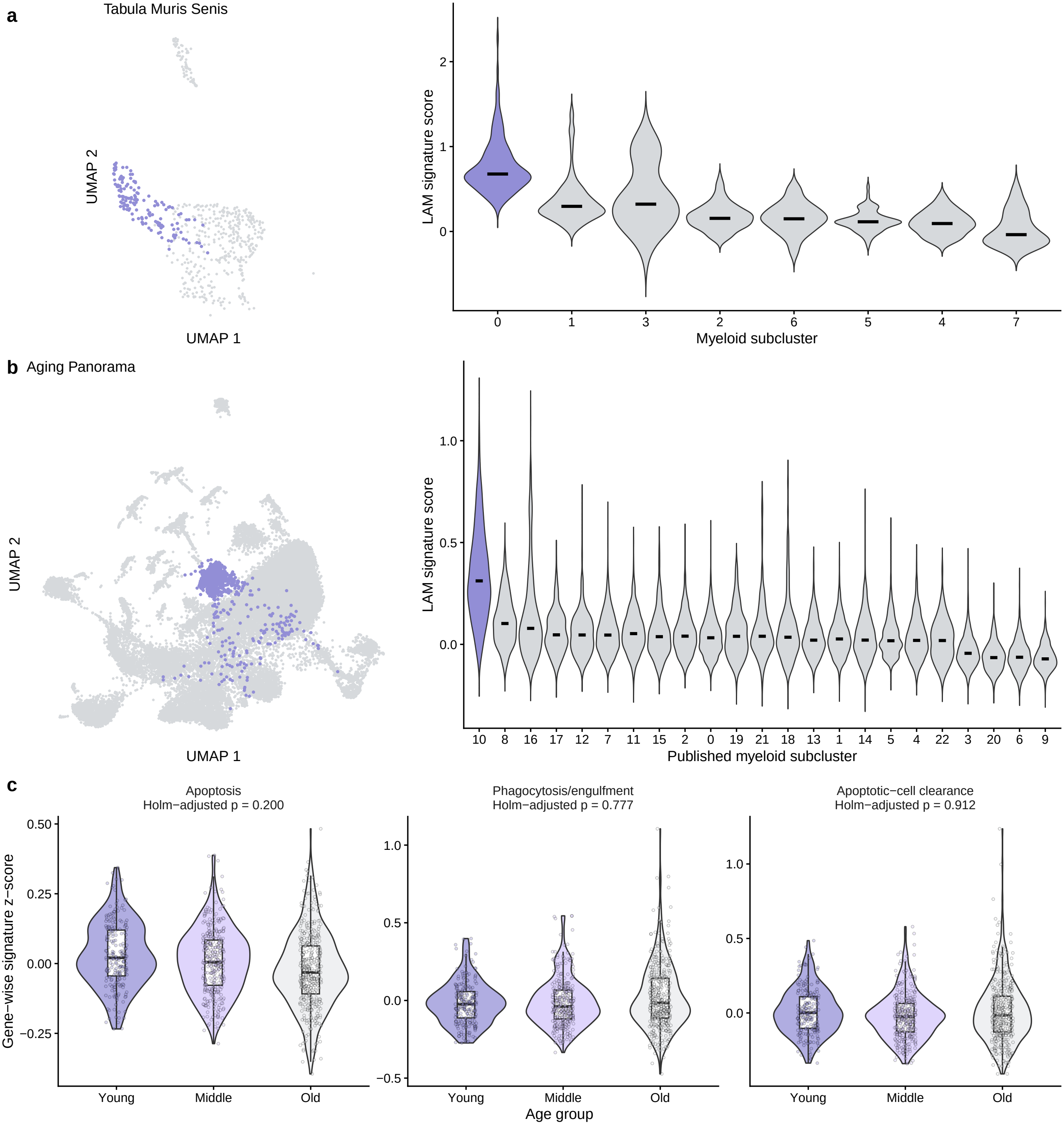


**Supplementary Figure 3**

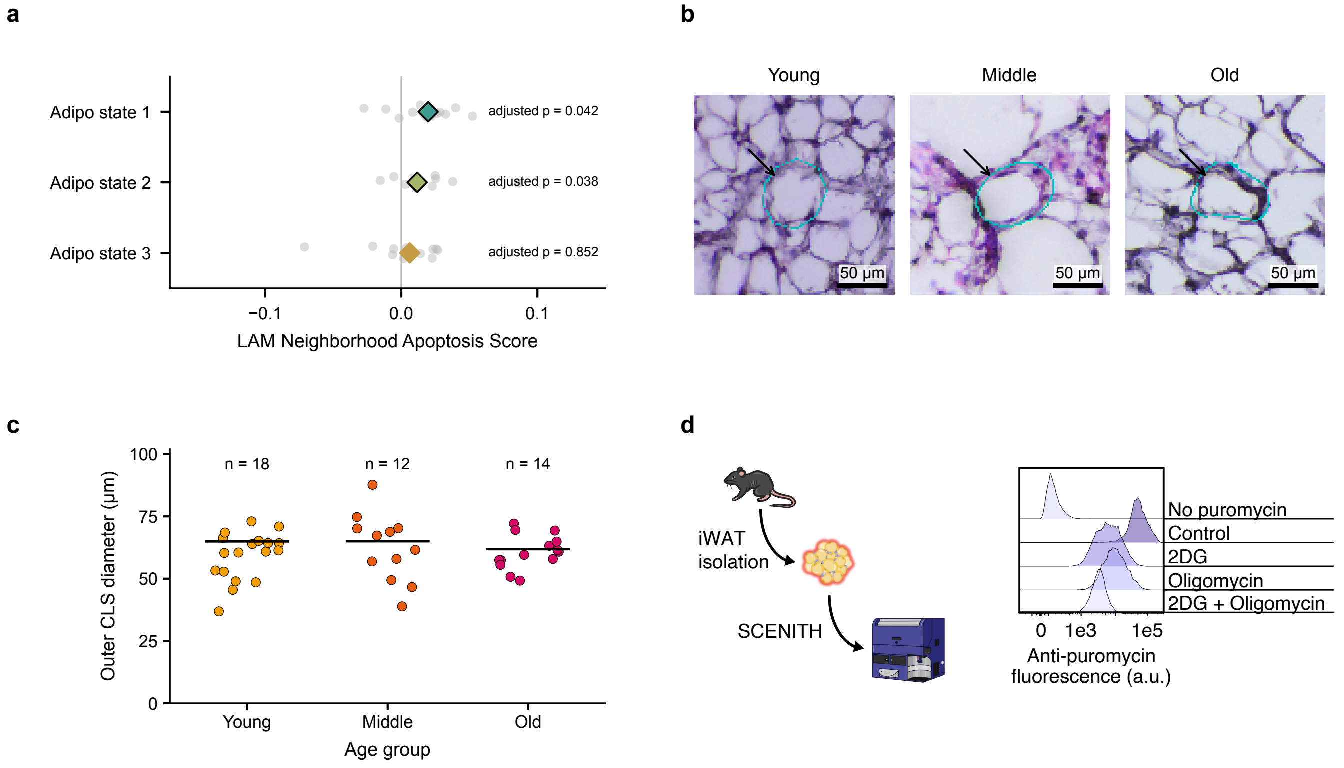


**Supplementary Figure 4**

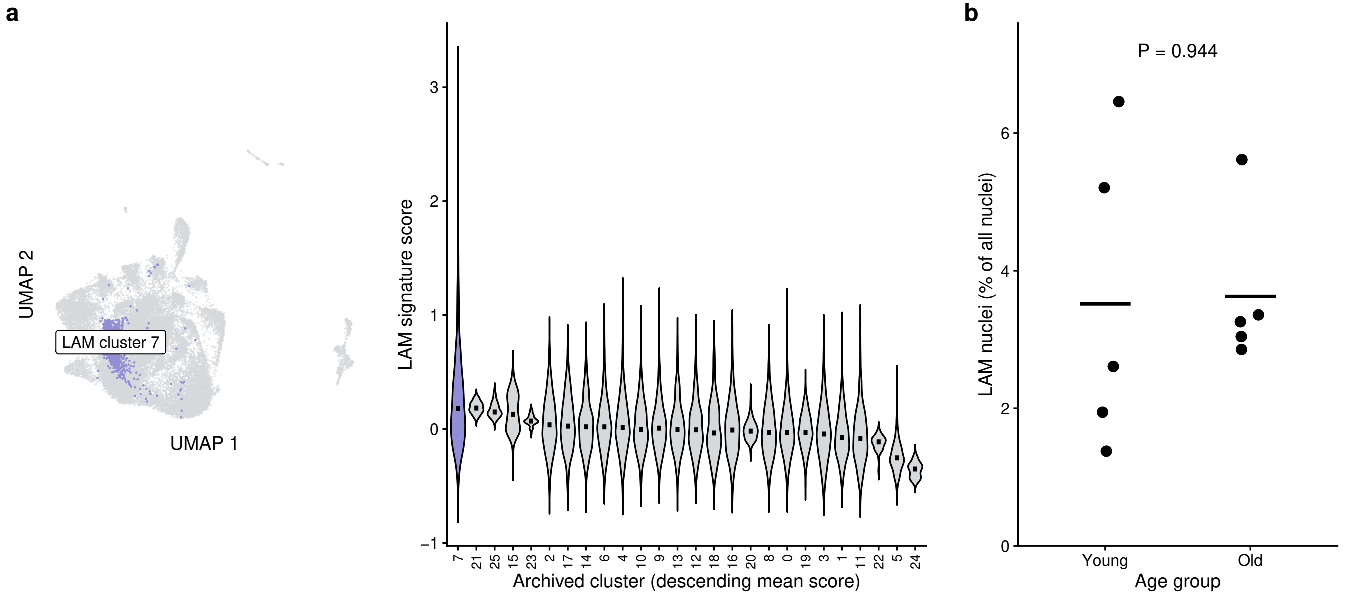


**Supplementary Figure 5**

### Supplementary Figure Legends

#### Supplementary Figure 1

**Representative histology, spatial LAM-signature distribution and consistency with RCTD**

**a**, Haematoxylin-and-eosin overview images embedded in the Visium object for one representative section with complete matched histology from each age group (scale bars, 1 mm). **b**, Global percentile of the normalized area-under-the-curve score for the 58-gene LAM signature in the same sections. All sections share the same scale; each circle represents one Visium spot. **c**, Difference between the median mapped RCTD weight in spots assigned to each project annotation and the median in all other spots, calculated separately for each of 12 biological samples. Dots denote samples, colours denote age groups and diamonds denote medians across samples; the dashed line marks zero. The merged Adipo annotation was compared with the adipocyte reference class. AUCell, area-under-the-curve-based gene-set scoring; H&E, haematoxylin and eosin; LAM, lipid-associated macrophage; RCTD, robust cell type decomposition.

#### Supplementary Figure 2

**Adipo-state proportions and age-associated expression across spatial populations**

**a,** Fraction of merged Adipo spots assigned to each state in each biological sample. Adjusted P values summarize the age term from binomial models; n.s., not significant. **b,** Overlap of genes upregulated in Old versus Young across nine non-Adipo populations and three Adipo states. Each column shows genes upregulated in exactly the populations marked by filled dots. Upper bars give gene counts for the 21 largest intersections (including ties). Right-hand bars count all upregulated genes per population. **c,** MAST log2 fold-change distributions for Middle versus Young and Old versus Middle, displayed as in Fig. 2d. Positive values indicate higher expression in the older group.

#### Supplementary Figure 3

**LAM-signature enrichment in reference myeloid populations and LAM process scores across age**

**a**, Male Tabula Muris Senis myeloid cells shown in the existing adipose UMAP (left) and LAM-signature score distributions across myeloid subclusters (right). Cluster 0, the highest-median-score cluster at Louvain resolution 0.4, is highlighted in purple (147 male cells; 60/55/32 at 12/72/96 weeks, from 4/2/4 donor IDs).

**b**, Male Aging Panorama myeloid nuclei shown in the supplied sub-UMAP (left) and LAM-signature score distributions across all 23 published myeloid subclusters (right). Published cluster 10 is highlighted in purple (1,176 male nuclei from 16 donor IDs).

In **a**,**b**, grey denotes unselected myeloid populations. The original 59-gene LAM signature was scored as mean log-normalized signature expression minus expression-matched control-gene expression, using corrected Panorama gene-symbol assignments. Clusters are ordered by median score across both sexes; male distributions are displayed. Black crossbars indicate male medians. Violin widths are normalized and do not encode population size. Clustering and score ranking use both sexes; temporal analyses use only males.

**c**, Age distributions of gene-expression scores for Hallmark apoptosis, phagocytosis/engulfment and apoptotic-cell clearance in spatially annotated LAM spots from this study. Points are individual LAM spots (952 in total), violins show score distributions and boxplots indicate the median, interquartile range (IQR) and 1.5-IQR whiskers. Scores are means of gene-wise z-standardized log-normalized expression across the genes in each signature, with standardization performed across LAM spots. All twelve biological samples are included, four at every age group. Statistical tests use the mean score per biological sample, not individual spots as independent replicates. Two-sided linear age-trend permutation tests enumerate all 34,650 age-label assignments retaining four samples per age group. P-values are jointly Holm-adjusted across the three signatures.

#### Supplementary Figure 4

##### Adipocyte state neighbourhoods and SCENITH analyses

**a**, Apoptosis-neighbourhood differences for Adipo states 1-3, calculated as in Fig. 4a, with BH correction across three states. **b,** Exemplary H&E images of histologically confirmed crown-like structures (CLS) in inguinal white adipose tissue from young, middle and old mice (arrows; scale bars, 50 µm). Cyan outlines delineate the central adipocyte-like space and its immediately surrounding cellular crown. **c**, Jitterplot of outer CLS equivalent-circle diameters. Each point represents one CLS, horizontal bars indicate the mean of mouse-specific medians. **d**, Schematic of inguinal white adipose tissue (iWAT) isolation from aging mice and subsequent SCENITH analysis by flow cytometry, alongside exemplary histograms of anti-puromycin fluorescence. Distributions show the background fluorescence in absence of puromycin, control condition without inhibitors, 2-deoxyglucose (2DG), oligomycin, and combined 2DG plus oligomycin treatment.

#### Supplementary Figure 5

##### Human LAM identity and donor representation

**a**, UMAP of the GSE235 529 single-nucleus RNA-sequencing dataset highlighting LAM cluster 7 (purple; other clusters, grey), and distributions of the project LAM signature score across all published clusters. The signature comprises 58 mapped human orthologues present in the SCT assay. Expression was standardized per gene across nuclei and averaged per nucleus. Violin plots show score distributions, with black crossbars indicating medians; clusters are ordered by decreasing mean signature score.

**b**, LAM nuclei as a percentage of all retained nuclei per male donor in GSE235 529. Points represent individual donors (n = 5 younger and 5 older); horizontal bars indicate group means.

### Supplementary Tables

#### Supplementary Table 1

**Information on processed samples of this study.**

#### Supplementary Table 2

**Age-associated differential gene expression across spatially annotated populations.**

#### Supplementary Table 3

**Gene overlap among late-age LAM transcriptional clusters.**
